# Limiting DNA polymerase delta alters replication dynamics and leads to a dependence on checkpoint activation and recombination-mediated DNA repair

**DOI:** 10.1101/2019.12.17.879544

**Authors:** Natasha C Koussa, Duncan J. Smith

## Abstract

DNA polymerase delta (Pol *δ*) plays several essential roles in eukaryotic DNA replication and repair. At the replication fork, Pol *δ* is responsible for the synthesis and processing of the lagging-strand. At replication origins, Pol *δ* has been proposed to initiate leading-strand synthesis by extending the first Okazaki fragment. Destabilizing mutations in human Pol *δ* subunits cause replication stress and syndromic immunodeficiency. Analogously, reduced levels of Pol *δ* in *Saccharomyces cerevisiae* lead to pervasive genome instability. Here, we analyze how the depletion of Pol *δ* impacts replication origin firing and lagging-strand synthesis during replication elongation *in vivo* in *S. cerevisiae.* By analyzing nascent lagging-strand products, we observe a genome-wide change in both the establishment and progression of replication. S-phase progression is slowed in Pol *δ* depletion, with both globally reduced origin firing and slower replication progression. We find that no polymerase other than Pol *δ* is capable of synthesizing a substantial amount of lagging-strand DNA, even when Pol *δ* is severely limiting. We also characterize the impact of impaired lagging-strand synthesis on genome integrity and find increased ssDNA and DNA damage when Pol *δ* is limiting; these defects lead to a strict dependence on checkpoint signaling and resection-mediated repair pathways for cellular viability.

**SIGNIFICANCE STATEMENT:** DNA replication in eukaryotes is carried out by the replisome – a multi-subunit complex comprising the enzymatic activities required to generate two intact daughter DNA strands. DNA polymerase delta (Pol *δ*) is a multi-functional replisome enzyme responsible for synthesis and processing of the lagging-strand. Mutations in Pol *δ* cause a variety of human diseases: for example, destabilizing mutations lead to immunodeficiency. We titrate the concentration of Pol *δ* in budding yeast – a simple model eukaryote with conserved DNA replication machinery. We characterize several replication defects associated with Pol *δ* scarcity. The defects we observe provide insight into how destabilizing Pol *δ* mutations lead to genome instability.

## INTRODUCTION

DNA polymerase delta (Pol *δ*) is an essential replisome component in all known eukaryotes (1). During lagging-strand synthesis, Pol *δ* extends Okazaki fragment primers synthesized by DNA polymerase alpha/primase (Pol α) (2). Pol *δ* plays an additional role at replication origins, synthesizing a stretch of DNA on the nascent leading strand that is subsequently extended by DNA polymerase epsilon (Pol ε) (3–6). Thus, Pol *δ* is directly responsible for the synthesis of approximately half the nuclear DNA in eukaryotic genomes (7,8) and is intimately involved in every step of the replication program. Multiple suppression mechanisms exist to maintain the specificity of Pol ε and Pol *δ* for leading- and lagging-strand synthesis, respectively (9,10). However, Pol *δ* can effectively synthesize the entire leading strand in both budding and fission yeast when the catalytic activity of Pol ε is abrogated (4, 11–14), and recent work suggests that Pol *δ* may take over leading-strand synthesis from Pol ε during replication termination under normal conditions (15).

Mutations affecting the stability or catalytic activity of Pol *δ* are associated with various human diseases. Alleles with reduced replication fidelity, including exonuclease-deficient alleles, are driver mutations for heritable cancers (16–18) and can induce catastrophic genome instability when expressed in yeast (19). A heterozygous active site mutation that abolishes Pol *δ* catalytic activity gives rise to developmental defects including lipodystrophy (20). In addition, mutations that reduce the stability of the Pol *δ* complex have recently been reported as causal for a syndromic immunodeficiency associated with replication stress (21). Some of the phenotypes associated with Pol *δ* deficiency overlap with those reported for Pol ε hypomorphy in a mouse model (22).

The depletion of Pol *δ* in budding yeast via transcriptional repression of the catalytic Pol3 subunit leads to various manifestations of genome instability, including an increase in the frequency of both point mutations and gross chromosomal rearrangements (23–25). However, the underlying basis for these defects could in theory derive from any of the roles of Pol *δ* in DNA replication or repair (26). The DNA replication machinery is highly conserved from yeast to humans. Therefore, directly characterizing the behavior of the yeast replisome under conditions where Pol *δ* is scarce can provide insights into the mechanisms by which Pol *δ* hypomorphs lead to genome instability in multicellular eukaryotes including humans.

Here, we define the short-term effects of both moderate and acute Pol *δ* depletion on the DNA replication program in budding yeast via the genome-wide analysis of lagging-strand synthesis, replication-origin firing and DNA polymerase usage. We find that even in severely limiting Pol *δ* conditions, no other replicative polymerase or translesion polymerase significantly contributes to lagging-strand synthesis. Pol *δ* depletion impairs the firing of all replication origins and slows lagging-strand replication. Even a slight reduction in Pol *δ* levels leads to increased DNA damage, which is only tolerated when resection-mediated repair pathways are intact. These genomic insults lead to a dependence on checkpoint activation for survival through even a single cell cycle.

## RESULTS

In *S. cerevisiae*, Pol *δ* is a heterotrimeric complex (27), comprising the essential Pol3 catalytic subunit and two additional accessory subunits – Pol31 and Pol32. Of the two accessory subunits, only Pol31 is essential for viability as long as the PIP boxes on the other subunits are intact (28). Besides forming part of Pol *δ*, Pol31 and Pol32 are integral components of the translesion DNA polymerase ζ (29): we therefore chose to titrate Pol3 to avoid as best as possible any indirect effects due to Pol ζ depletion. Titratable expression of *POL3* from the *GAL1* promoter has previously been used to study the long-term effects of Pol *δ* depletion (23,24). However, because modulating transcription using galactose is relatively slow and *GAL1-POL3* strains have been reported to revert to high expression with detectable frequency (24), we chose to titrate Pol3 via targeted proteolysis. The resulting rapid degradation of Pol3 allows us to investigate the immediate consequences of Pol *δ* depletion within a single cell cycle.

### Titration of Pol *δ in vivo* via controlled proteolysis of Pol3

To allow the rapid depletion of Pol3 from cycling *S. cerevisiae* cells, we tagged the protein with an C-terminal auxin-inducible degron (30) with an additional C-terminal 9xMyc tag for detection of Pol3 by western blot. Pol3 depletion in a strain carrying TIR1 (30) was readily detectable after 30 minutes (**Fig. S1A**) and treatment with indoleacetic acid (IAA) at concentrations between 0.01 mM and 1 mM allowed Pol3 levels to be titrated between wild-type and essentially undetectable levels (**Fig. 1A**).

**Figure 1.**
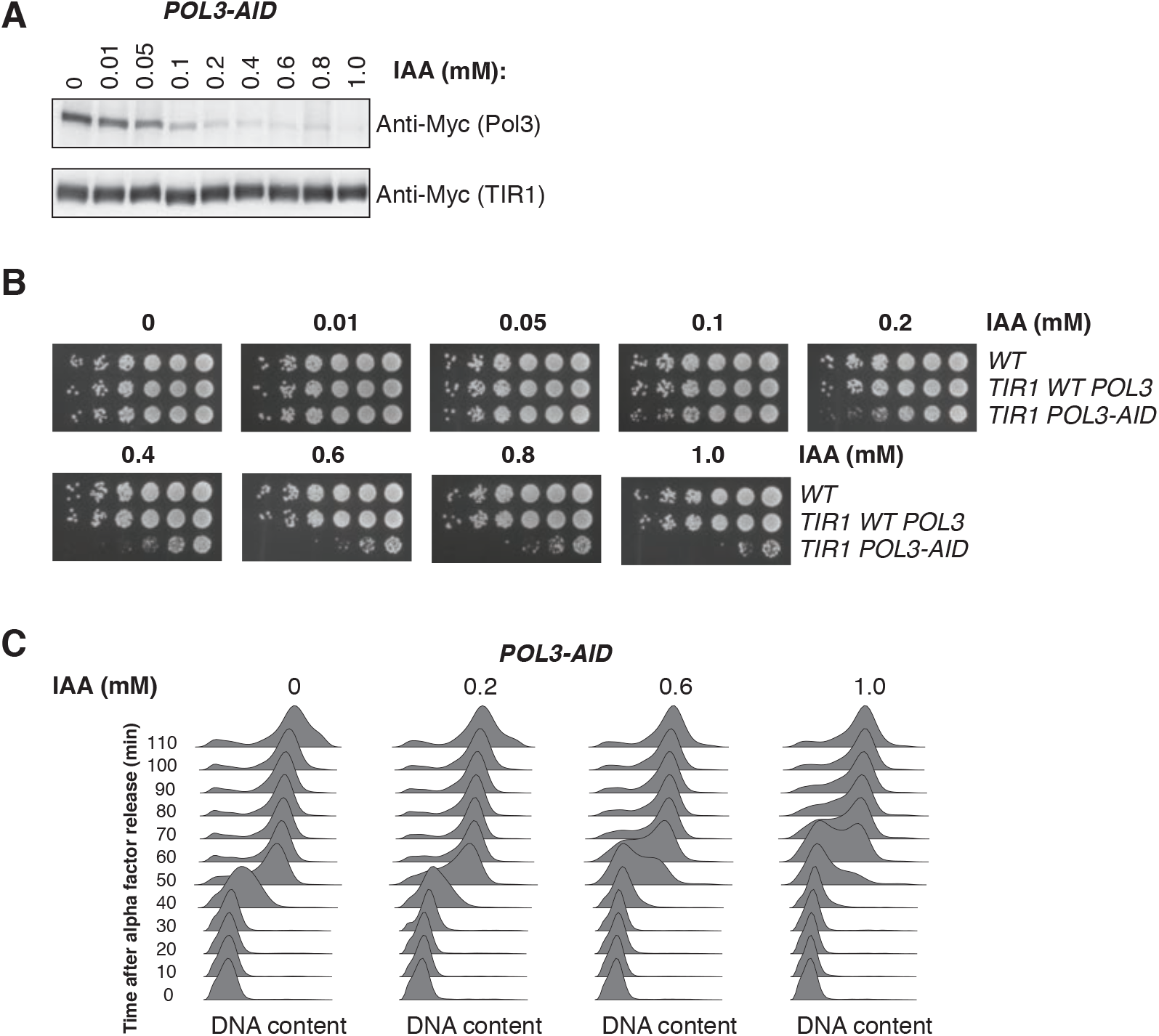
Titration of Pol3 *in vivo* delays and extends S-phase, leading to slow growth. **A.** Western blot against Pol3-9Myc in asynchronous cultures of the *POL3-AID* strain following 2h of treatment with the indicated concentration of IAA. Myc-tagged TIR1 is required for degradation of Pol3, and serves as a loading control. **B.** Serial dilution spot tests of *POL3-AID* cells and two control strains on YPD plates supplemented with IAA at concentrations from 0 to 1 mM. **C.** DNA content measured by flow cytometry for *POL3-AID* cells released from α-factor-mediated G1 arrest at 30°C. Individual cultures were treated with the indicated concentration of IAA for 2h during the initial arrest, and subsequently released into media containing the same concentration of IAA.

Long-term growth of *POL3-AID* cells was impaired in the presence of IAA when assayed by serial dilution of cells on solid media (**Fig. 1B**) or cell density in liquid media (**Fig. S1B**). However, *POL3-AID* cells remained viable even at 1 mM IAA – the highest concentration we tested (**Fig. 1B, S1B**). Analysis of DNA content by flow cytometry in synchronized cultures indicated that low Pol3 levels both delayed entry into S-phase and extended its duration (**Fig. 1C**). Consistent with these data, asynchronous cultures treated with high concentrations of IAA accumulated in G1 and S-phase (**Fig. S1C**).

### Effects of limiting Pol *δ* on productive replication origin firing *in vivo*

Pol *δ* is responsible for the majority of DNA replication on the lagging-strand (7,8), but also plays a key role in the initiation of leading-strand synthesis at replication origins (3–5). Pol *δ* depletion led to slower entry into S-phase (**Fig. 1C**), suggesting that origin firing may be impaired in limiting Pol *δ* conditions. To investigate the effect of Pol3 depletion on origin firing, we sequenced Okazaki fragments from strains depleted for Pol3. As previously described (31), the Origin Efficiency Metric (OEM) can be determined from Okazaki fragment sequencing based on the fraction of Okazaki fragments mapping to either the Watson or Crick strand in a window ±10 kb around previously validated replication origins: OEM represents the fraction of cells in the population in which a given origin fires. However, when comparing OEM between samples it is necessary to correct for differences in cell-cycle stage across the population. As shown schematically in **Fig. S2A**, the OEM for all but the earliest-firing, most efficient origins will change throughout S-phase due to an increasing contribution of passive replication from forks established at neighboring origins. A moderately efficient origin will first appear to have high origin efficiency because the only Okazaki fragments around the origin result from its stochastic early firing in a fraction of cells in the population. As S-phase progresses, cells in which this origin did not fire replicate this region via replisomes emanating from other origins, resulting in a decrease in Okazaki fragment strand bias – and a lower OEM – around the origin (**Fig. S2A**).

Because Pol3 depletion alters the cell-cycle distribution of the population (**Fig. 1C, S1C**), we used synchronized cells to determine the effects low Pol *δ* levels on origin firing. Alpha-factor arrested cultures were treated with rapamycin to deplete DNA ligase I (Cdc9) from the nucleus via anchor-away (32) and released into S-phase in YPD alone or YPD supplemented with 1 mM IAA. Rapamycin and IAA were maintained throughout the release and cells were sampled every five minutes for both flow cytometry (**Fig. S2B**) and Okazaki fragment sequencing. We confirmed that the OEMs decrease as replication proceeds, even without Pol3 depletion (**Fig. S2A&C**). In order to compare origin efficiency between cells with endogenous Pol3 levels and the more slowly progressing Pol3 depletion conditions, we monitored the extent of replication at each timepoint by plotting normalized Okazaki fragment coverage for both cells grown in YPD alone or supplemented with 1mM IAA across all sequenced timepoints (**Fig. S2D&E**). As expected, at early timepoints, coverage is variable across the genome: high coverage correlates with genomic locations with early origins. At later timepoints, coverage in later replicating regions increases as S-phase proceeds. In order to directly compare origin efficiency, different timepoints were selected for 0mM and 1mM IAA conditions such that the genome coverage is similar between the two conditions. We therefore chose two sets of timepoints, one in early S and one in mid S phase, in order to compare origin efficiency (**Fig. S2F&G**).

Origin efficiency was significantly reduced at both early and mid S timepoints in Pol3 depletion conditions (**Fig. 2A**) (paired t-test, p-value < 0.0001). In order to determine if both efficient and inefficient origins were impacted to the same degree, we directly compared individual origin efficiencies with and without Pol3 depletion. Regardless of the origin efficiency in endogenous conditions, Pol3 depletion led to lower origin efficiency in both early S (**Fig. 2B**) and mid S timepoints (**Fig. 2C**). As expected, lowered origin efficiency in Pol3 depletion led to a lower strand bias for Okazaki fragments around origins in early S (**Fig. 2D**) and mid S timepoints (**Fig. 2E**). By calculating the change in strand bias in Pol3 depletion conditions and comparing between early and mid S timepoints, we show a similar change in strand bias in both timepoints (**Fig. 2F**). Taken together, we conclude that limiting Pol *δ* lowers origin firing globally without differentially impacting early- or late-firing replication origins.

**Figure 2.**
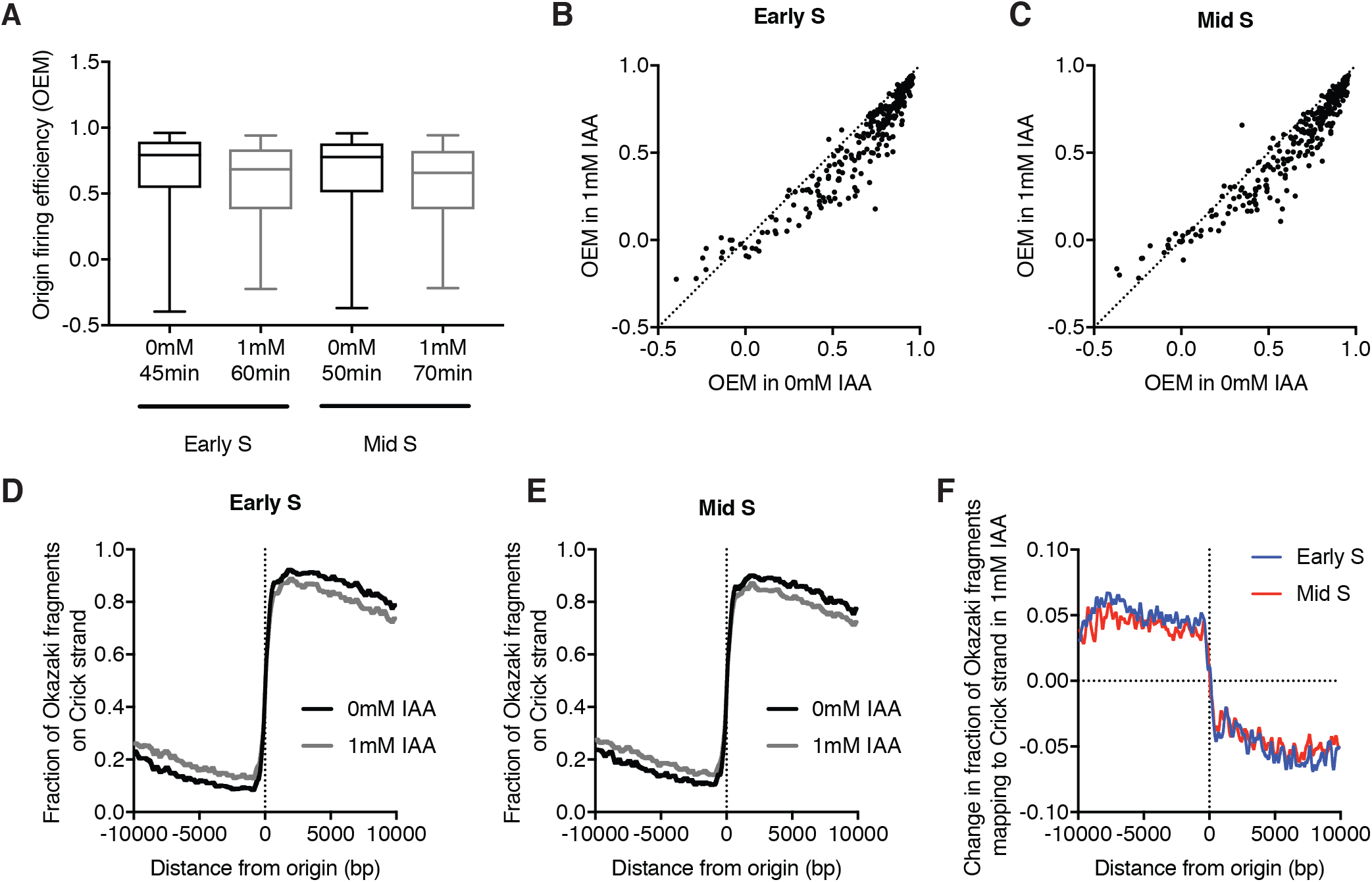
Pol3 depletion decreases firing efficiency of replication origins. **A.** Replication-origin firing efficiencies of synchronous *POL3-AID* cells in YPD ±1 mM IAA (in addition to rapamycin treatment to deplete nuclear Cdc9 by anchor away). Origin Efficiency Metrics (OEMs) were calculated from Okazaki fragment distributions around 281 high-confidence origins (31) for early S or mid S timepoints. Data represent the mean efficiency and whiskers indicate minimum and maximum. **B-C.** Replication-origin firing efficiency (OEMs) in *POL3-AID* cells in YPD ±1 mM IAA, for early S (B) or mid S (C) timepoints (see text and Fig. S2 for details of how early- and mid-S were defined). The line of identity is shown as a dotted line. **D-E.** Okazaki fragment strand bias around origins, calculated as the fraction of total reads mapping to the Crick strand (representing rightward-moving replication forks). Data from cells grown in YPD are shown in black, YPD + 1 mM IAA is shown in grey for early S (D) or mid S (E) timepoints. **F.** Change in Okazaki fragment strand bias around origins, calculated by subtracting the data obtained with 1mM IAA from 0mM IAA.

### Alternative polymerases do not significantly contribute to lagging-strand synthesis when Pol *δ* is limiting

Pol *δ* is responsible for the bulk of lagging-strand synthesis (7). Previous work has shown that Pol *δ* can synthesize the leading strand when Pol ε is catalytically inactive. Yeast has three replicative polymerases (Pol α, Pol ε, Pol *δ*) and three translesion polymerases (4, 11–14). Pol ζ, Pol η, Rev1). Synthesis by Pol α is inhibited by RFC-dependent loading of PCNA (11,33), but Pol ε has been reported to compete weakly with Pol *δ* for lagging-strand primers *in vitro* (9). In addition, any of the three translesion polymerases present in *S. cerevisiae* could theoretically contribute to lagging-strand replication when Pol *δ* is limiting: indeed, Pol η has recently been reported to participate in lagging-strand synthesis to a measurable extent in unperturbed cells (34).

To test the possibility that translesion (TLS) DNA polymerases might contribute substantially to DNA replication during Pol3 depletion, we combined the *POL3-AID* allele with individual knockouts of *REV1, REV3* (Pol ζ), and *RAD30* (Pol η), as well as the *rev1 rev3 rad30* triple mutant (*ΔTLS*). Individual or simultaneous deletion of the three TLS polymerases failed to suppress or exacerbate the growth defect observed during Pol3 depletion at any concentration of IAA (**Fig. 3A**). We conclude that growth during transient Pol *δ* depletion is not rescued by the widespread action of TLS polymerases.

**Figure 3.**
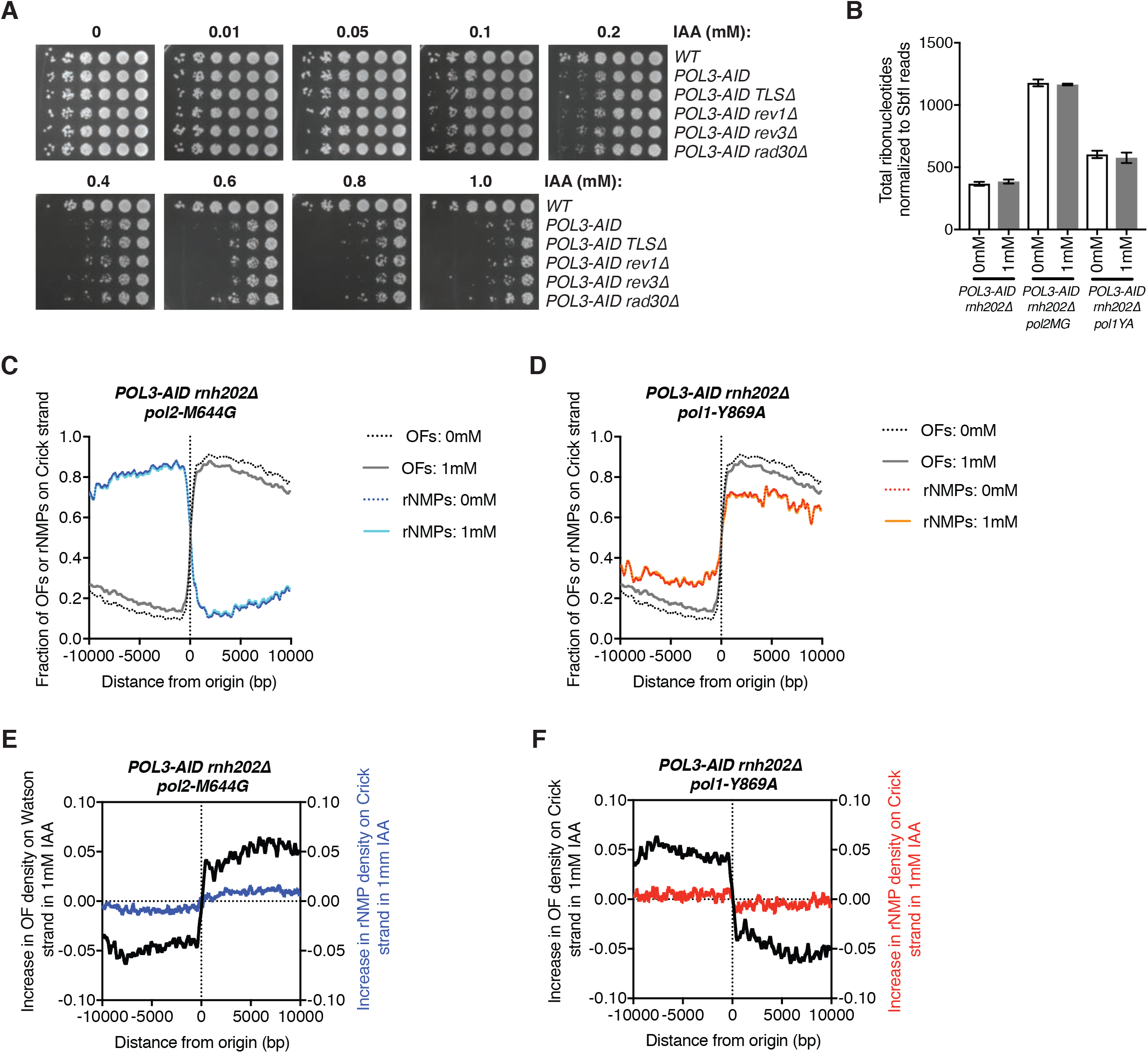
No significant use of alternative polymerases on the lagging-strand in Pol3 depletion. **A.** Serial dilution spot tests of *POL3-AID* cells with the indicated additional mutations on YPD plates supplemented with IAA at concentrations from 0 to 1 mM. **B.** Total number of ribonucleotides identified by HydEn-seq (7) normalized by reads mapping to SbfI sites. Asynchronous cultures of the indicated strains were treated ±1 mM IAA for 2h prior to cell collection. Data shown are the average of two biological replicates, with whiskers representing standard deviation. **C.** Fraction of ribonucleotides identified by HydEn-seq (blue) or Okazaki fragments (black/gray, data from Fig. 2) mapping to the Crick strand around the same set of replication origins analyzed in Fig. 2. Asynchronous cultures of the *POL3-AID pol2-M644G rnh202Δ* strain were treated ±1 mM IAA for 2h prior to cell collection. Data shown are the average of two biological replicates. Data for individual replicates are in Fig. S3. **D.** Fraction of ribonucleotides identified by HydEn-seq (red/orange) or Okazaki fragments (black/gray, data from Fig. 2) mapping to the Crick strand around the same set of replication origins analyzed in Fig. 2. Asynchronous cultures of the *POL3-AID pol1-Y869A rnh202Δ* strain were treated ±1 mM IAA for 2h prior to cell collection. Data shown are the average of two biological replicates. Data for individual replicates are in Fig. S3. **E.** Change in the fraction of ribonucleotides mapping to the Crick strand (blue) and Okazaki fragments mapping to the Watson strand (black, data from Fig. 2) around the same replication origins after 1mM IAA treatment in *POL3-AID pol2-M644G rnh202Δ* strains. The Okazaki fragment polarities are chosen such that the direction of the strand bias change inferred matches the strand bias of *pol2-M644G* ribonucleotide incorporation. **F.** Change in the fraction of ribonucleotides mapping to the Crick strand (red) and Okazaki fragments mapping to the Crick strand (black, data from Fig. 2) around the same replication origins after 1mM IAA treatment in *POL3-AID pol1-Y869A rnh202Δ* strains. The Okazaki fragment polarities are chosen such that the direction of the strand bias change inferred matches the strand bias of *pol1-Y869A* ribonucleotide incorporation.

In order to investigate the overall contribution of Pol ε to DNA replication in the context of Pol *δ* depletion, we combined *POL3-AID* with the *pol2-M644G* allele of the catalytic subunit of Pol ε: this allele causes increased incorporation of ribonucleotides into the nascent leading strand (35). We analogously used a *pol1-Y869A* allele to determine the contribution of Pol α to genome-wide DNA synthesis (7). In strains unable to remove single ribonucleotides from genomic DNA due to deletion of the RNase H2 subunit *RNH202*, ribonucleotide positions can be tracked by HydEn-seq: ribonucleotides are cleaved by alkaline hydrolysis, and the locations of the resulting nicks determined by strand-specific high-throughput sequencing (7). As previously described, we included a normalization step that effectively acts as a loading control by treating DNA with the rare-cutting SbfI restriction enzyme (4). We prepared HydEn-seq libraries from *POL3-AID rnh202Δ* strains, *POL3-AID pol2-M644G rnh202Δ* strains, and *POL3-AID pol1-Y869A rnh202Δ* strains grown ± 1 mM IAA for two hours before genomic DNA preparation. Two biological replicates were used for each condition. Average results for the two biological replicate strains at 0 mM and 1 mM IAA are shown in **Fig. 3**, and results for individual replicates in **Fig. S3**.

In order to determine whether the overall contribution of Pol ε or Pol α to the synthesis of nascent genomic DNA increases during Pol3 depletion, we quantified total ribonucleotides incorporated during Pol3 depletion, normalized to reads from SbfI sites. As expected, there were more ribonucleotides in the *POL3-AID pol2-M644G rnh202Δ* and *POL3-AID pol1-Y869A rnh202Δ* strains than *POL3-AID rnh202Δ* strains in 0mM IAA. If Pol ε or Pol α contributes more DNA synthesis during Pol3 depletion, we should observe an increase in ribonucleotide density relative to SbfI sites upon IAA treatment in *POL3-AID pol2-M644G rnh202Δ* or *POL3-AID pol1-Y869A rnh202Δ*, respectively. Notably, there was no significant increase in ribonucleotide density in either strain upon Pol3 depletion, consistent with neither Pol ε nor Pol α synthesizing a large amount of additional DNA during Pol3 depletion (**Fig. 3B**).

Ribonucleotides incorporated by Pol ε are normally highly specific for the leading strand, so an increased contribution of Pol ε to lagging-strand synthesis during Pol *δ* depletion should result in a loss of strand bias for Pol ε-derived ribonucleotides (**Fig. 6B**). On the other hand, ribonucleotides incorporated by Pol α normally show a bias towards the leading strand, due to the repeated priming of the lagging-strand; increased contribution of Pol α during lagging-strand synthesis should increase this strand bias. The strand bias of Pol ε (**Fig. 3C**) and Pol α (**Fig. 3D**) are very slightly shifted during Pol3 depletion. However, Pol3 depletion lowers the firing efficiencies of replication origins (**Fig. 2**), which means that the same strand cannot necessarily be assigned as lagging at different auxin concentrations, as seen by the strand bias of the Okazaki fragments. We therefore compared the change in ribonucleotide incorporation by Pol ε or Pol α to the change in lagging-strand bias inferred from the Okazaki fragment sequencing experiments shown in **Fig. 2**. Since the change in ribonucleotide strand bias in Pol3 depletion is less than the change in Okazaki fragment strand bias, we conclude that Pol ε and Pol α do not synthesize the lagging-strand to a significant extent, even in limiting Pol *δ* conditions. Together these data confirm that Pol *δ* is solely responsible for virtually all lagging-strand replication, even when Pol *δ* is severely limiting.

### Replication speed is reduced when Pol *δ* is limiting

The increased duration of S-phase that we observed under limiting Pol *δ* conditions (**Fig. 1C**) could be due to checkpoint activation and/or reduced replication speed. To directly test whether replication of the lagging-strand proceeds more slowly through the genome upon Pol3 depletion, we sequenced Okazaki fragments from strains synchronously released into S-phase at 25 °C in YPD ±1 mM IAA (flow cytometry data showing the timepoints selected for analysis are in **Fig. S2B**). Okazaki fragment sequencing data from this time course are shown in **Fig. 4A**. The density of sequencing reads in cultures sampled from 45-60 minutes after release showed decreased replication of origin-distal regions in the samples in which Pol3 was depleted (**Fig. 4A**), consistent with reduced replication speed.

**Figure 4.**
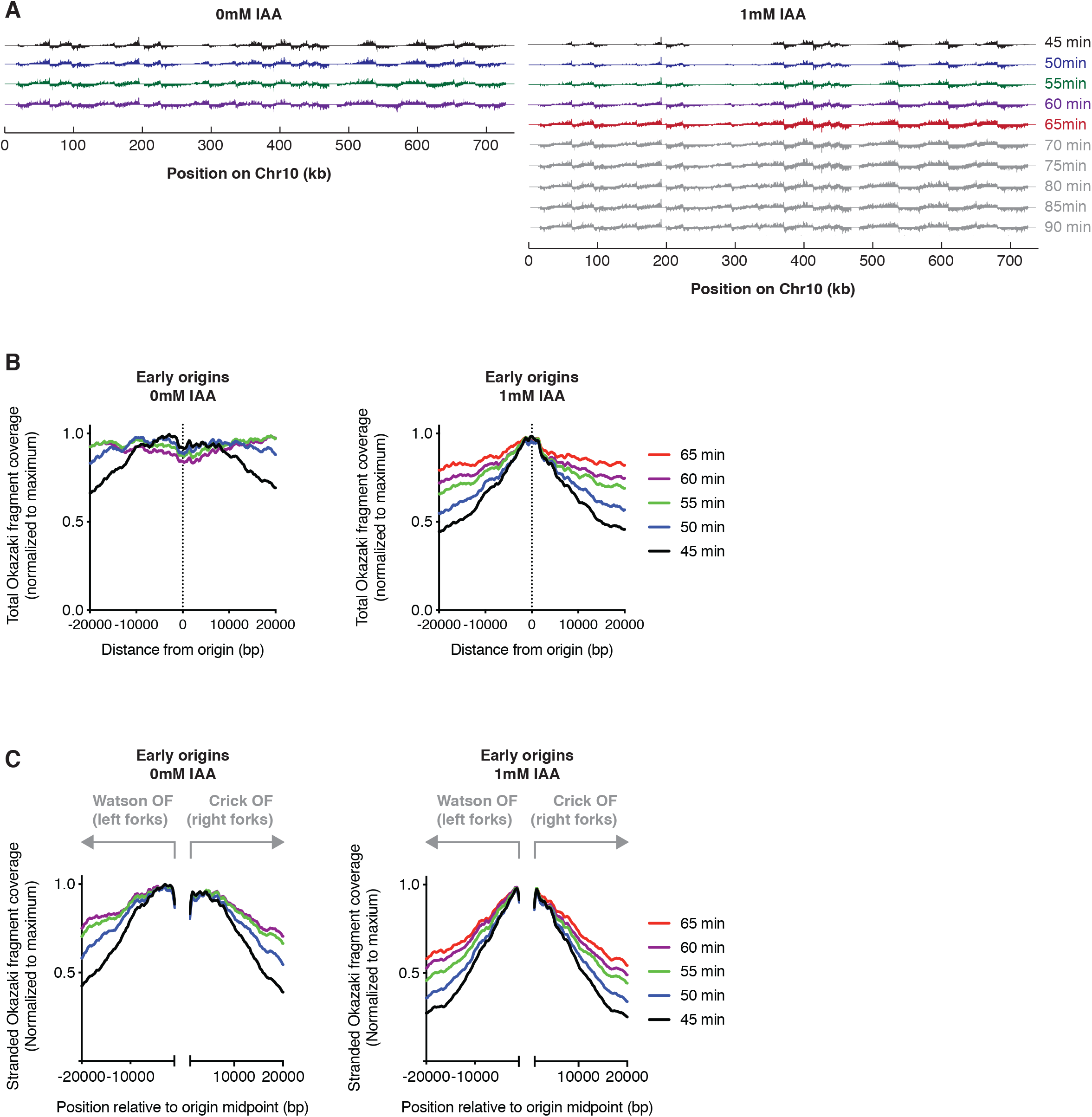
Pol3 depletion slows replication fork progression. **A.** Okazaki fragment sequencing data across chromosome 10 for cultures released from α-factor-mediated G1 arrest at 25°C. IAA treatment during and following G1 arrest was maintained as in Fig. 1C and 2D. Data are displayed such that Watson strand hits are above the axis and Crick strand hits below the axis for each timepoint. Timepoints shown in color are analyzed in B and C. Flow cytometry data for this experiment are in Fig. S3 **B.** Total coverage of Okazaki fragments on both the Watson and Crick strands across a 40 kb region around early-firing origins (defined as in Fig. 3D) at the indicated time and concentration of IAA. Data are shown with a 100 bp bin size, and are normalized to the maximum signal in the range such that complete replication of the regions ±20 kb from these origins will result in a flat line at 1.0. **C.** Coverage of Watson-strand Okazaki fragments in the region from −20 kb to −1 kb, and Crick-strand Okazaki fragments in the region from +1 kb to +20 kb around the early-firing origins analyzed in B. This analysis specifically measures the progression of leftward- and rightward-moving forks emanating from these origins. As in B, data are normalized to the maximum signal in the entire 40 kb range.

To confirm that Pol3 depletion globally decreases replication speed, we carried out meta-analysis of both total (**Fig. 4B**) and strand-specific (**Fig. 4C**) Okazaki fragment density 20 kb up- and downstream of early-firing replication origins. Total Okazaki fragment density monitors the extent to which the lagging-strand has been replicated, regardless of the direction of fork movement. By contrast, the stranded analysis considers only Okazaki fragments from leftward-moving forks to the left of the origin, and from rightward-moving forks to the right – these data therefore report more directly on the progression of replication away from the origins being analyzed. In both cases, Okazaki fragments from Pol3-depleted cells generated a sharper peak close to early-firing origins after 45 minutes, which propagated away from origins more slowly than observed in samples from cells grown in YPD (**Fig. 4B, 4C**). We therefore conclude that the progression of replication through the genome is impaired when Pol *δ* is limiting. Our analysis cannot distinguish between repeated stalling of replication and reduced speed or replisome movement.

### Conditional depletion of Pol *δ* leads to accumulation of RPA and dependence on checkpoint activation

Limiting levels of Pol *δ* could lead to gaps in the daughter strand and/or excess single-stranded DNA at the replication fork. To explore the possibility that single-stranded DNA is more abundant when Pol *δ* is scarce, we assayed the level of RPA bound to chromatin via western blot against Rfa1 following subcellular fractionation (**Fig. 5A**). After 2 hours of IAA treatment, both the total and chromatin-bound pools of RPA increased substantially. To exclude the possibility that this increase simply reflected the accumulation of IAA-treated cells in S-phase (**Fig. S1C**), we assayed Rfa1 levels after synchronous release into S-phase (**Fig. 5B**). Chromatin-bound RPA was highly enriched in both mid and late S-phase in IAA-treated cells compared to untreated controls, implying the persistence of single-stranded DNA throughout S-phase.

**Figure 5.**
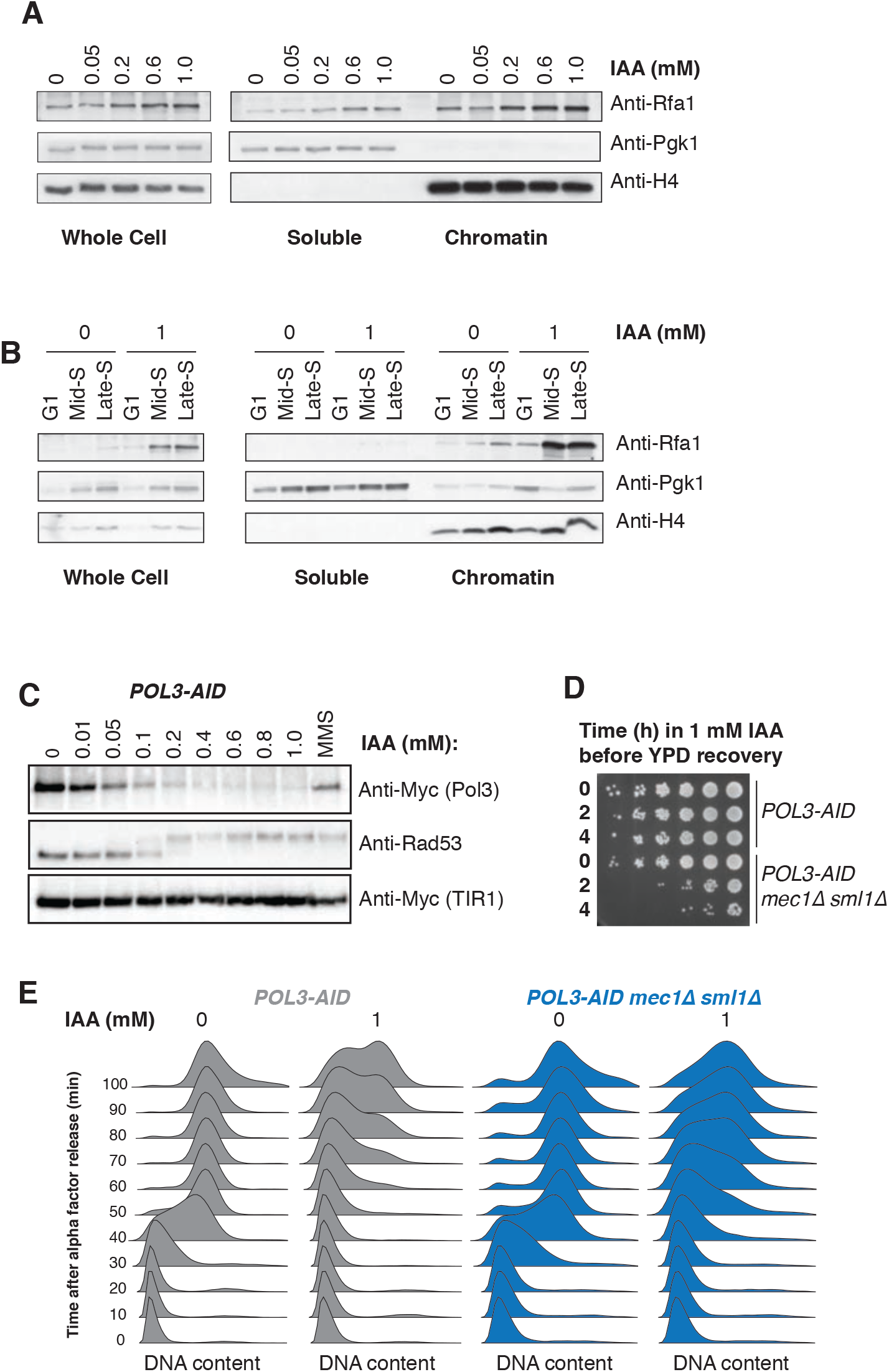
Pol3 depletion leads to RPA accumulation on chromatin and checkpoint activation. **A.** Western blot against Rfa1 on soluble and chromatin-bound fractions of log-phase *POL3-AID* cells treated with the indicated IAA concentration for 2h. **B.** Western blots as in (A), but using cells collected during G1 arrest, or in mid- or late S-phase as indicated. **C.** Western blots to detect Rad53 phosphorylation in asynchronous cultures of *POL3-AID* cells grown in YPD supplemented with IAA. **D.** Serial dilution spot tests of *POL3-AID* or *POL3-AID mec1Δ sml1Δ* cells exposed to 1 mM IAA for 0, 2 or 4h of growth during logarithmic phase, followed by plating on YPD without IAA. **E.** DNA content measured by flow cytometry for *POL3-AID* or *POL3-AID mec1Δ sml1Δ* cells released from α-factor-mediated G1 arrest at 30°C. As in Fig. 1C, individual cultures were treated with the indicated concentration of IAA for 2h during the initial arrest, and subsequently released into media containing the same concentration of IAA.

Chromatin-bound RPA can activate the S-phase checkpoint, therefore we assayed checkpoint activation during Pol3 depletion by western blot against Rad53 following 2 hours of treatment with increasing concentrations of IAA (**Fig. 5C**). Phosphorylation of Rad53 was apparent from 0.2 mM IAA – the lowest concentration that leads to a growth defect in *POL3-AID* strains (**Fig. 1B**). In order to determine if this checkpoint activation response is necessary for survival in limiting Pol3 conditions, we assayed viability by serial dilution spot tests of cells with or without the apical kinase *MEC1,* allowed to recover on YPD plates after transient treatment in liquid culture with 1mM IAA (**Fig. 5D**). Depletion of Pol3 for 2-4 hours was sufficient to cause widespread cell death in the *mec1Δ sml1Δ* background. Under these conditions, G1-arrested *mec1Δ sml1Δ* cells depleted for Pol3 were able to enter S-phase, but few if any cells reached G2 100 minutes after release from G1 (**Fig. 5E**). We conclude that the checkpoint is required to survive even a single cell cycle when Pol ∂ is severely limiting.

### Conditional depletion of Pol *δ* leads to DNA damage and dependence on recombination-mediated repair

Rad53 phosphorylation can occur via the DNA replication checkpoint (DRC), dependent on Mrc1, or the DNA damage checkpoint (DDC), mediated by Rad9 (36). We tested Rad53 phosphorylation in *POL3-AID* strains deleted for either *MRC1* or *RAD9*, or for the apical kinase *MEC1*, which is required for both pathways (**Fig. 6A**). As expected, Rad53 phosphorylation was virtually abolished in a *mec1Δ sml1Δ* strain. Deletion of *RAD9* substantially reduced the extent of Rad53 phosphorylation, especially at lower concentrations of IAA, while deletion of *MRC1* had a much weaker effect. Furthermore, consistent with the relative contributions of the DDC and DRC to Rad53 phosphorylation (**Fig. 6A**), a *POL3-AID rad9Δ* strain had a stronger growth defect than a *POL3-AID mrc1Δ* strain at moderate concentrations of IAA (**Fig. 6B**). We therefore conclude that Pol *δ* scarcity leads to checkpoint activation predominantly through the DDC, but with a contribution from the DRC.

**Figure 6.**
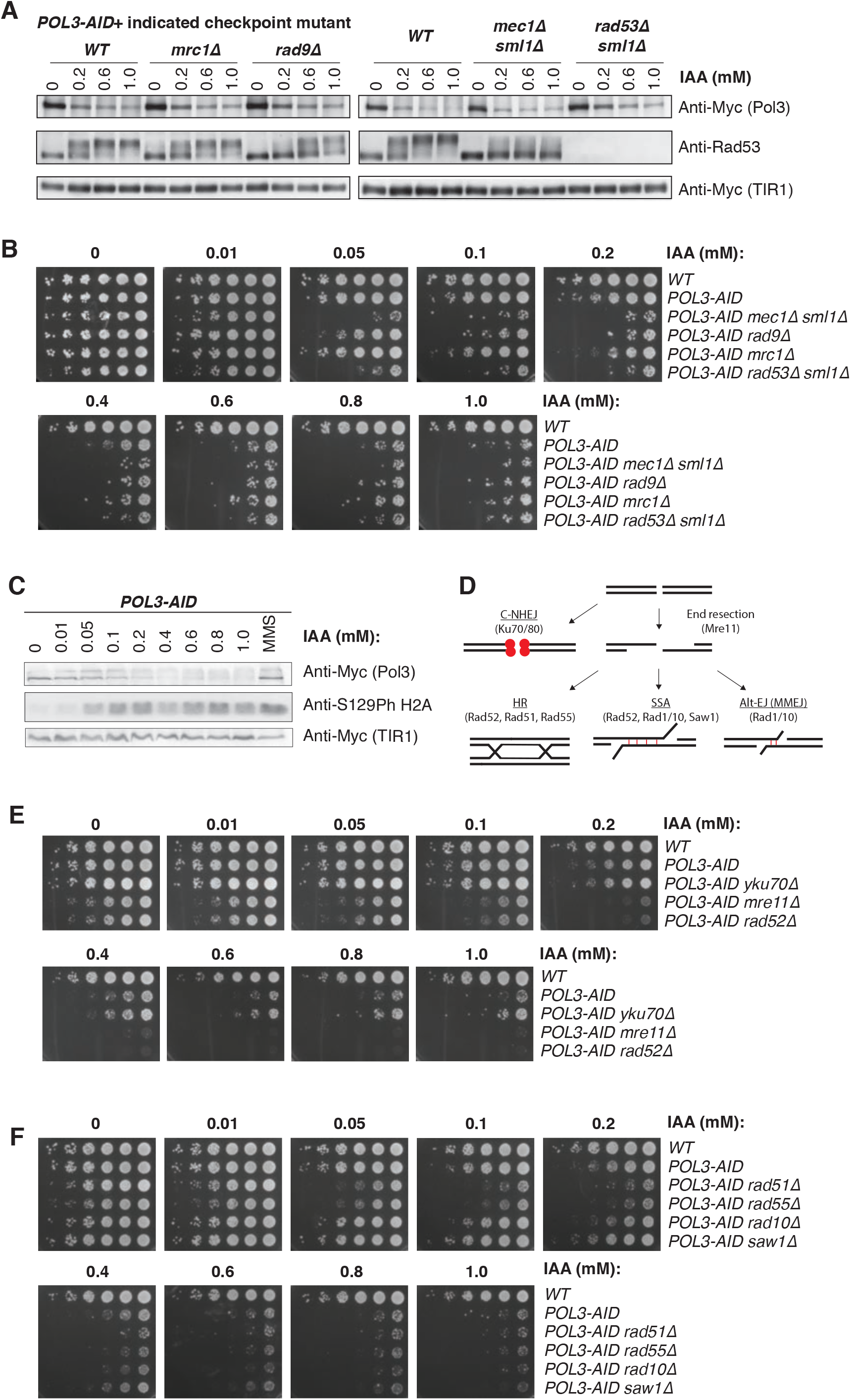
Pol3 depletion leads to DNA damage and dependence on homologous recombination. **A.** Western blots to detect Rad53 phosphorylation in asynchronous cultures of the indicated strains grown in YPD supplemented with IAA. **B.** Serial dilution spot tests of *POL3-AID* cells with the indicated additional mutations on YPD plates supplemented with IAA at concentrations from 0 to 1 mM. **C.** Western blot to detect histone H2A phosphorylation at S129 in asynchronous cultures of the *POL3-AID* strain grown in YPD supplemented with IAA as indicated. **D.** Schematic of major repair mechanisms with key factors deleted in Figure 6E&F indicated. **E.** Serial dilution spot tests of *POL3-AID* cells with the indicated additional mutations on YPD plates supplemented with IAA at concentrations from 0 to 1 mM. **F.** Serial dilution spot tests of *POL3-AID* cells with the indicated additional mutations on YPD plates supplemented with IAA at concentrations from 0 to 1 mM.

The requirement for checkpoint activation, particularly via the DDC, suggested that DNA damage may accumulate during Pol3 depletion. We tested for the accumulation of DNA double strand breaks (DSBs) via western blots against histone H2A phosphorylated at Serine 129 (phosphor-H2A) following 2 hours of IAA treatment (**Fig. 6C**) (37). Phospho-H2A was detectable at IAA concentrations over 0.05 mM – the same concentration at which *mec1Δ sml1Δ* cells begin to show an obvious growth phenotype (**Fig. 6B&C**), suggesting that DSBs arise during even moderate depletion of Pol ∂. DSBs can be repaired by multiple repair mechanisms, principally non-homologous end joining (NHEJ) or resection-based mechanisms including homologous recombination (HR), single strand annealing (SSA), and alternate end joining (alt-EJ) (**Fig. 6D**) (38). To test which pathway is required for the repair of DSBs caused by Pol3 depletion, we first tested if survival was dependent on NHEJ by deleting *YKU70*, or by one of the end-resection based mechanisms by deleting the nuclease component of the MRX complex, *MRE11* or the mediator of recombination, *RAD52* (39,40). Deletion of *RAD52* or *MRE11* substantially decreased viability at moderate IAA concentrations; by contrast, abrogation of end-joining via deletion of *YKU70* slightly increased fitness under these conditions (**Fig. 6E**).

In order to determine which end-resection based mechanisms are responsible for repair during Pol3 depletion, we combined deletion of *RAD51, RAD55* (both essential for HR), *SAW1* (essential for SSA), or *RAD10* (essential for SSA and alt-EJ) (38, 41–44) with the *POL3-AID* allele. Deletion of *RAD51* or *RAD55* most severely impacted viability of cells at moderate IAA concentration, indicating that HR is primarily responsible for repair of DSBs during Pol3 depletion (**Fig. 6F**). Together, these data suggest that depletion of Pol3, and consequently Pol *δ*, rapidly leads to the accumulation of single-stranded DNA and double-strand breaks, leading to checkpoint activation and delayed progression through S-phase. HR-mediated DNA repair pathways are essential for cells with limiting Pol *δ* to complete S-phase and continue through the cell cycle.

## DISCUSSION

By directly analyzing the products of lagging-strand synthesis following proteolytic titration of Pol3 in *S. cerevisiae*, we identified several immediate consequences of Pol *δ* depletion for DNA replication. Despite a strict requirement for Pol *δ* for synthesis of the lagging-strand (**Fig. 3**), extremely low levels of Pol *δ* support viability if the checkpoint and resection-mediated repair pathways are intact (**Fig. 5 & 6**). Both replication origin firing (**Fig. 2**) and the progression of replication through the genome (**Fig. 4**) are impaired by reduced levels of Pol *δ*.

We find that limiting levels of Pol *δ* cause damage during replication (**Fig. 6C)**, which lead to a dependence on resection-based repair mechanisms for survival, particularly HR (**Fig. 6E&F**). The presence of DSBs and dependence on HR for repair may explain the high incidence of large indels in diploid *S. cerevisiae* strains grown in low Pol *δ* conditions (24,25). DSBs can arise from replication-associated ssDNA gaps (45) and breakpoints of LOH events in diploid strains with low levels of Pol *δ* were found to correlate with locations of ssDNA detected by APOBEC-induced mutagenesis (46).

Indeed, we detect higher levels of chromatin-bound RPA during Pol *δ* depletion, indicative of ssDNA (**Fig. 5A&B**). Similarly, a recent report in human cells detected higher levels of RPA-bound ssDNA upon Pol *δ* knockdown (47). While Pol *δ* hypomorphs in humans show evidence of underreplicated DNA and DSBs, it is unknown if repair of damage due to hypomorphic Pol *δ* is repaired by HR or another mechanism in humans (21).

Under normal conditions, Pol *δ* is solely responsible for bulk lagging-strand synthesis. This division of labor is favorable due to the high error rate of Pol α (48) and the inability of Pol ε to carry out the strand-displacement synthesis required for Okazaki fragment processing (49). The exclusion of other polymerases from the lagging-strand is mediated by RFC. Loading of PCNA by RFC at the primer-template junction recruits Pol *δ* and is thought to inhibit extensive synthesis or rebinding of Pol α, preventing bulk synthesis of the lagging-strand by this error-prone polymerase (50). RFC outcompetes Pol ε for the primer terminus thereby excluding Pol ε from the lagging-strand while CMG protects Pol ε from RFC-mediated inhibition on the leading strand (14,50). Our data demonstrate that, even when Pol ∂ levels are dramatically reduced, other polymerases do not contribute substantially to lagging-strand synthesis. (**Fig. 3**). Therefore, it appears that the RFC-based suppression mechanisms that enforce lagging-strand synthesis by Pol *δ* are robust enough to prevent either Pol α or Pol ε from extending a significant fraction of primers even when Pol *δ* is extremely scarce. Previous work has found evidence of synthesis by Pol η primarily on the lagging-strand. However, it remains unclear whether Pol η has a direct role during replication of the lagging-strand, or whether the observed Pol η activity is due to PCNA-mediated recruitment of Pol η to endogenous damage on the lagging-strand (34). While it is possible that TLS polymerases may be more active on the lagging-strand during Pol *δ* depletion, their contribution is, at most, minor as deletion of any or all of these polymerases do not affect growth during Pol *δ* depletion.

Although traditional models of lagging-strand synthesis imply the recruitment of a new Pol *δ* for the synthesis of each Okazaki fragment, recent work in *S. cerevisiae* has shown that a single Pol *δ* can synthesize multiple Okazaki fragments without dissociating, both *in vitro* and *in vivo* (6,51). Indeed, the *in vivo* residence time of Pol *δ* on chromatin may be sufficiently long to allow a single Pol *δ* to synthesize the lagging-strand throughout the entire lifetime of a replisome from initiation to termination (51). *In vitro* single-molecule data suggest that Pol *δ* may undergo concentration-dependent exchange, such that high concentrations of the complex result in exchange while lower concentrations lead to stable association of Pol *δ* with the replisome (6). These data cannot easily be reconciled with the effects we observe upon extreme Pol *δ* depletion. During mid-S-phase in a wild-type *S. cerevisiae* cell, there are an estimated 300 replisomes (52) and approximately 3000 Pol *δ* complexes (53).

Although our Western blots are not strictly quantitative, we are confident that the higher IAA concentrations used in our experiments deplete Pol3 by more than 90% (**Fig. 1A & S1A**), such that there should be fewer Pol *δ* complexes than the normal number of active replisomes. While we cannot formally exclude the possibility that Pol3 depletion might be uneven across the population, we do not observe the presence of distinct populations when we assay S-phase progression by flow cytometry (**Fig. 1C**).

Intuitively, severe depletion of a polymerase that remains stably associated with the replication fork would be expected to allow the establishment of normal replisomes at early-firing origins, with later-firing origins unable to fire – at least until the early replisomes had encountered a convergent replication fork and terminated. Those replisomes established before Pol *δ* was exhausted would also, according to this model, have a normal complement of replicative polymerases and therefore be expected to move at a normal speed without leaving gaps or exposing large amounts of single-stranded DNA at the replication fork. In contrast to these predictions, we observe a relatively minor decrease in replication-origin firing that affects all origins approximately equally (**Fig. 2),** along with defects in replication that include slower lagging-strand synthesis (**Fig. 4**), increased RPA association with chromatin (**Fig. 5A&B**), and a strict dependence on the checkpoint (**Fig. 5D**). Furthermore, the overall timing profile of replication, while slower upon Pol *δ* depletion, is remarkably consistent across a huge variation in Pol *δ* concentration (**Fig. S2D&E**).

One possible explanation for our data could be the presence of a population of replisomes associating with Pol *δ*, and another population completely lacking Pol *δ*. *In vitro*, leading strand synthesis can proceed without Pol delta (5, 10,11). Replisomes lacking Pol *δ* would leave one daughter strand unreplicated, activating the checkpoint. Another possibility is that a single Pol *δ* could be shared between two or more replisomes – either in the context of associated sister replisomes or a replication factory (54–56). Reducing the average number of Pol *δ* complexes at a cluster of two or more associated replisomes could lead to competition between substrates for Pol ∂, leading to checkpoint activation and slower replisome progression.

## MATERIALS AND METHODS

### Yeast strains

All strains are W303 RAD5+ background and contained additional mutations required for anchor-away depletion of Cdc9. The genotype of the wild-type strain is *tor1-1::HIS3, fpr1::NatMX4, RPL13A-2xFKBP12::TRP1, CDC9-FRB::HygMX*. The Pol3 depletion strain contains OsTIR1 integrated at the *URA3* locus and *POL3* tagged with IAA17(71-116)-9xmyc (Addgene #99522) as described in (57). Individual knockout strains were created by gene replacement. Ribonucleotide-hyperincorporating strains were a generous gift from Hannah Klein. All mutations were introduced into the *POL3-AID* background by cross, except for *rad52*. Biological replicate strains represent independent colonies derived from one or more crosses.

### Cell growth, spot tests, and cell-cycle synchronization

Yeast were grown in YPD at 30C unless indicated otherwise. 3-indoleacetic acid (IAA) (Sigma I2886-5G) was dissolved in 95% ethanol to 200mM. For short term experiments, IAA was added to log phase cells for two hours, followed by addition of 1ug/mL rapamycin (Spectrum 41810000-2) for one hour for Okazaki fragment analysis. Spot tests were performed with exponentially growing cultures (OD 0.65), washed in sterile water, and diluted five-fold in sterile water in 96-well plates and spotted on plates at 30C for two days. For synchronized S-phase analysis, log phase cells (OD 0.2) were briefly sonicated to disrupt clumps and 10ug/mL alpha factor was added, followed by 5ug/mL every hour until >95% cells were fully shmooed. After complete arrest, IAA was added and 5ug/mL alpha factor was added every hour to maintain arrest. To release cells from arrest, cells were washed twice with deionized water and resuspended in YPD with or without IAA as required.

### Flow cytometry

Cells were collected by adding 150uL of yeast culture to 350uL absolute ethanol and stored at 4C. Samples were treated with RNase by pelleting cells and resuspending in 500uL of 50mM sodium citrate with 42ug/mL RNase A and incubating at 50C for two hours, followed by addition of 100ug proteinase K for two additional hours. Equal volume of 50mM sodium citrate with 0.2uL SYTOX green (Fisher S7020) was added, samples were sonicated, and analyzed on a Becton Dickinson Accuri.

### Subcellular fractionation and western blotting

Cells for subcellular fractionation were treated with 0.1% sodium azide and collected by centrifugation and stored at −20C. Cells were thawed on ice, washed and resuspended in spheroplasting buffer, and treated with 10uL of 20mg/mL zymolyase T20 and 1uL 1M DTT for 40 minutes at 30C before harvesting spheroplasts by centrifugation. Spheroplasts were resuspended in 300uL extraction buffer and divided into whole cell (50uL), soluble (50uL), and chromatin (200uL) fractions. The whole cell fraction was treated with 1.25uL 10% Triton-100, vortexed, incubated on ice for 5 minutes, treated with 1uL universal nuclease, incubated for 15 minutes on ice, and mixed with 20uL urea loading buffer. The soluble fraction was treated with 1.25uL 10% Triton-100, vortexed, incubated on ice for 5 minutes, centrifuged, and 20uL urea loading buffer was added to the supernatant. The chromatin fraction was treated with 1.25uL 10% Triton-100, vortexed and incubated on ice for 5 minutes before addition of 30% sucrose. After centrifugation, the previous step was repeated, and the resulting pellet was resuspended in 50μL EB with 1.25μL 10% Triton-100, treated with 1uL universal nuclease, incubated for 15 minutes on ice, and mixed with 20uL urea loading buffer.

Cells for western blotting were collected (2.0 OD units), washed, and resuspended in 200uL 20% TCA with glass beads, vortexed for 10 minutes, and the lysate was kept. The beads were washed with 600uL 5% TCA, combining with the lysate from the previous step. The lysate was centrifuged and the pellet was resuspended in 100uL urea loading buffer with 1uL 10M NaOH to restore color.

All western blotting samples were boiled for five minutes at 95C before loading onto an SDS-PAGE gel. Gels were blotted onto nitrocellulose membranes by wet transfer, blocked in 5% milk in PBS-T for one hour, and incubated with primary antibody at 4C overnight. After PBS-T washes, secondary antibody incubation for one hour, and PBS-T washes, blots were exposed with SuperSignal West Pico (Fisher 34080). The following antibodies were used in this study: anti-myc (1:1000 Genscript A00173-100), anti-Rad53 (1:1000 Abcam ab104232), anti-S129phosp H2A (1:1000 Abcam ab15083), anti-Rfa1 (1:50000 Abcam ab221198), anti-H4 (1:1000 Abcam ab10158), anti-PGK1 (1:1000 Fisher 459250), anti-rabbit IgG HRP (1:5000 GE Healthcare NA934V), anti-mouse IgG HRP (1:5000 Abcam ab97046).

### Okazaki fragment preparation, labeling and sequencing

Cells were grown as described above for a synchronous time course. Okazaki fragments were purified, end-labeled and deep-sequenced as previously described (58,59), with minor modifications. Briefly, DNA was purified and Okazaki fragments were purified using sequential elutions from Source 15Q columns and treated with RNase before adaptor ligation, second strand synthesis, and barcoding. Paired-end sequencing (2 × 75 bp) was carried out on an Illumina Next-seq 500 platform.

### HydEN-seq

HydEN-seq was carried out as described (4). Briefly, DNA was cut with SbfI-HF, treated with 300mM KOH, and 5’ phosphorylated before adapter ligation, second strand synthesis and barcoding. Paired-end sequencing (2 × 75 bp) was carried out on an Illumina Next-seq 500 platform.

### Computational analyses

FASTQ files were aligned to the S288C reference genome (SGD, R64-2-1) using the Bowtie (v2.2.9). Low quality reads and PCR duplicates were removed and resulting data was converted to BEDPE files using the Samtools suite (v1.3.1). For Okazaki fragment sequencing, genome coverage was calculated using the Bedtools suite (v2.26.0) in a strand-specific manner. Origin efficiency metric analysis of predefined origins was carried out as previously described (31) with the origin list from the same source (included as table S1). Briefly, the fraction of Okazaki fragments mapping to the Watson or Crick strand in 10 kb windows to the left and right of the origin (W_L_, W_R_, C_L_, and C_R_) is calculated for each origin. OEM is calculated as W_L_/(W_L_+C_L_) – W_R_/(W_R_+C_R_). For HydEN-seq, reads were aligned to the genome as above and duplicates were removed. The location of ribonucleotides were determined as in (7) and locations mapping to SbfI sites were removed. Total ribonucleotides reads were determined by summing all ribonucleotides on both strands for all chromosomes except for chromosome 12 (as this contains the rDNA repeats) and divided by the total reads at SbfI sites (for all chromosomes except 12). For origin analysis around previously described origins, Crick reads were divided by total reads at each location around origins and plotted. Calculations were done in R with custom in-house scripts.

## DATA AVAILABILITY

Sequencing data have been submitted to the GEO under accession number 141884.

## ACKNOWLEDGEMENTS

We thank Tom Petes and Dirk Remus for helpful discussions. We thank Hannah Klein for sharing strains. We thank members of the Smith lab for discussions and critical reading of the manuscript, and NYU Gencore for assistance with Illumina sequencing. This work was supported by NIH R01 GM114340 and R35 GM134918 to D.J.S.

## LEGENDS TO SUPPLEMENTARY FIGURES

**Supplementary figure 1(associated with figure 1).**
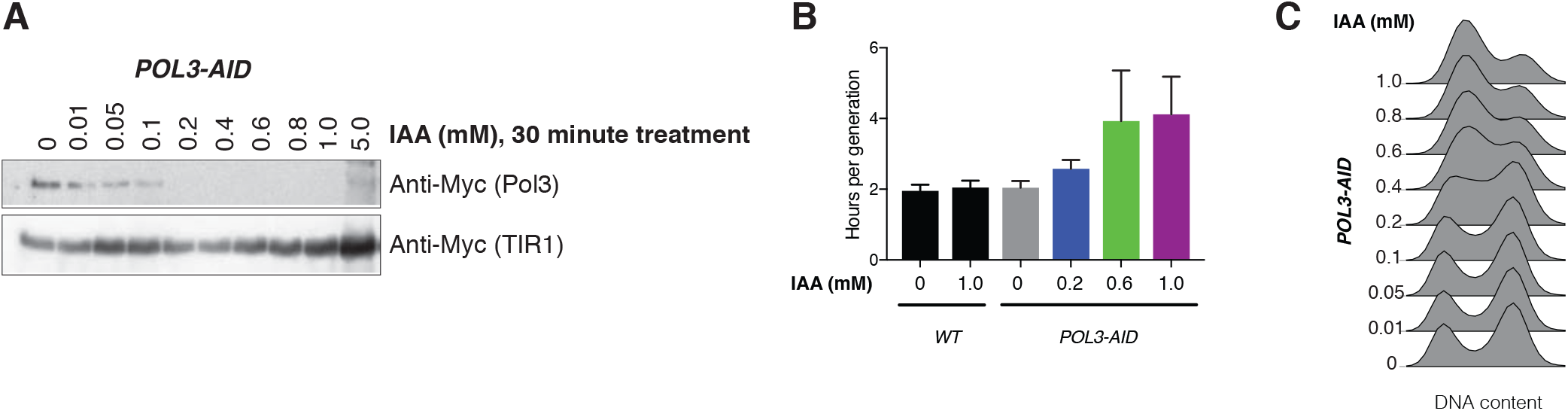
Further characterization of Pol3 depletion kinetics and the effect of depletion on growth rate. **A.** Western blot against Pol3-9Myc or OsTIR1-9Myc in asynchronous cultures of the *POL3-AID* strain following 30 minutes of treatment with the indicated concentration of IAA. **B.** Growth rates of *POL3-AID* cells in liquid culture. Data were calculated from three replicates. **C.** DNA content measured by flow cytometry for logarithmically growing *POL3-AID* cells treated with the indicated concentration of IAA for 2h.

**Supplementary figure 2 (associated with figure 2).**
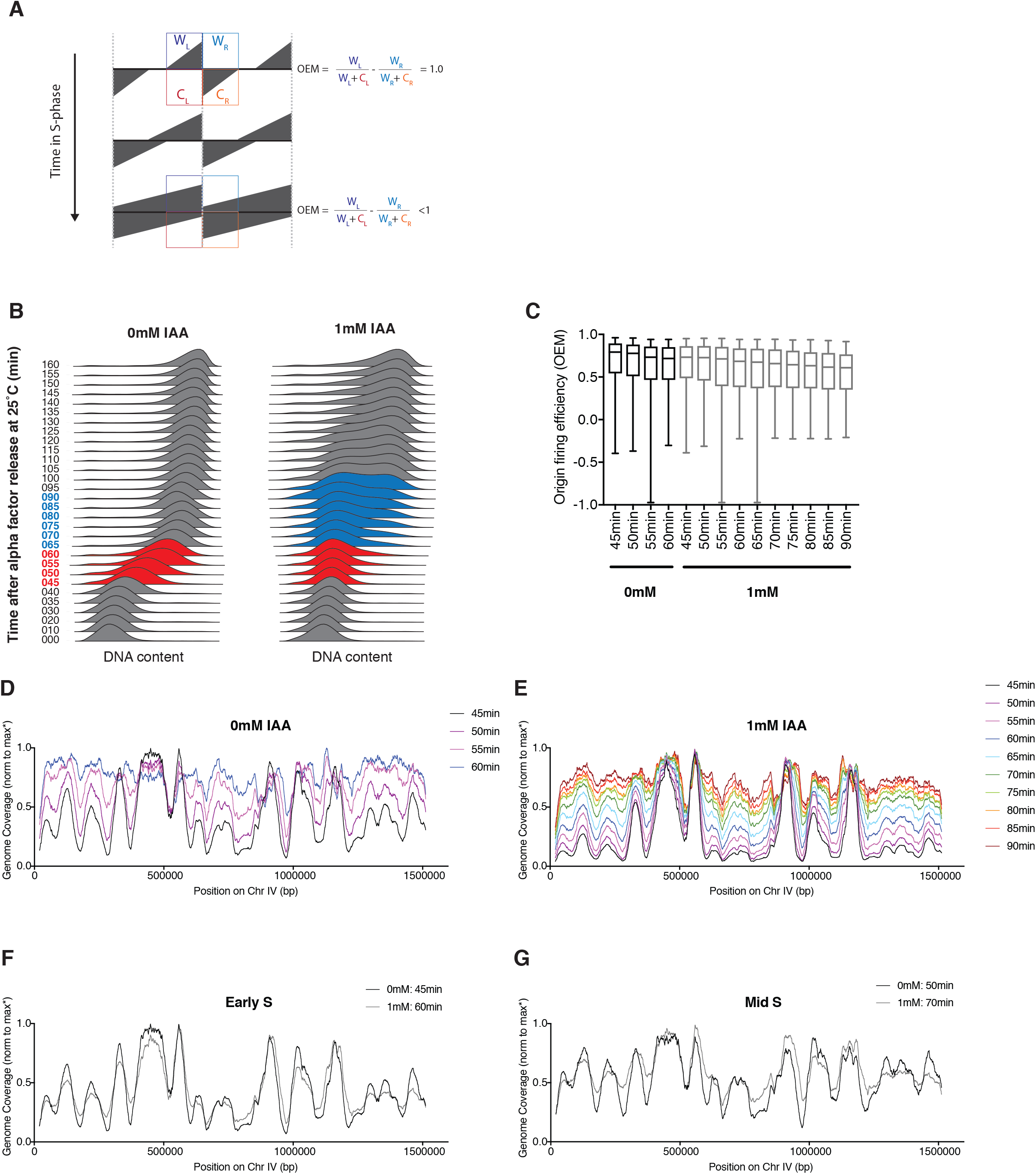
**A.** Schematic of expected Okazaki fragment distributions and OEMs for a moderately efficient replication origin (as calculated in (31)) over the course of S-phase. Okazaki fragments (grey) emanate from origins (dashed lines) to the left on the Watson strand (WL) or to the right on the Crick strand (CR). As S-phases progresses, the middle origin is passively replicated by the forks from the origins on the left and right in a fraction of cell, resulting in a lower OEM. **B.** Samples used for analysis of origin firing (Figure 2) and replication speed (Figure 4). DNA content measured by flow cytometry for the samples shown in Fig. 4A. Red timepoints were sequenced for both 0 and 1 mM IAA, and blue timepoints for 1 mM only. **C.** Replication-origin firing efficiency for samples sequenced from Supp Figure 2A, calculated as OEM from Okazaki fragment distributions around 281 high-confidence origins (31) for each sequenced time point across S phase. Data represent the mean efficiency averaged across two replicate strains at each time point. Whiskers indicate minimum and maximum. **D-E.** Total coverage of Okazaki fragment sequencing data across chromosome 4 for synchronous *POL3-AID* cultures treated with the indicated concentration of IAA for 2h and rapamycin for 1h treatment to deplete Cdc9 from the nucleus by anchor away. Data are normalized to the maximum of non-repetitive regions. **F-G.** Total coverage of Okazaki fragment sequencing data across chromosome 4 as in Supp Figure 2C-D for matched timepoints in early (F) or mid (G) S phase.

**Supplementary figure 3 (associated with figure 3).**
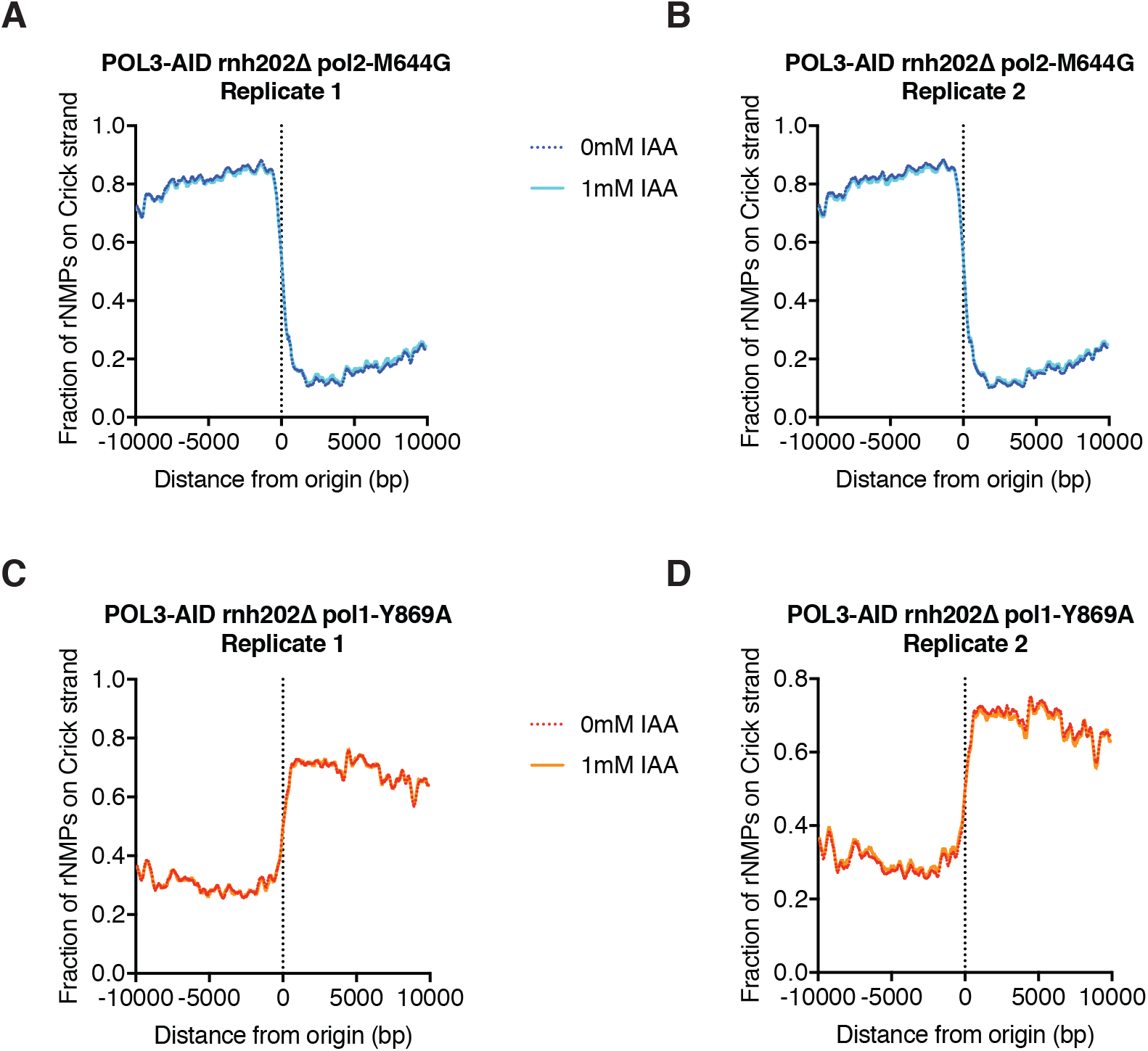
**A-B.** Analysis of ribonucleotide distribution around replication origins in the *POL3-AID pol2- M644G rnh202Δ* genetic background as shown for pooled replicates in Fig. 3C, separated by individual biological replicate strain as indicated. **C-D.** Analysis of ribonucleotide distribution around replication origins in the *POL3-AID pol1- Y869A rnh202Δ* genetic background as shown for pooled replicates in Fig. 3D, separated by individual biological replicate strain as indicated.

**Supplementary figure 4 (associated with figure 5).**
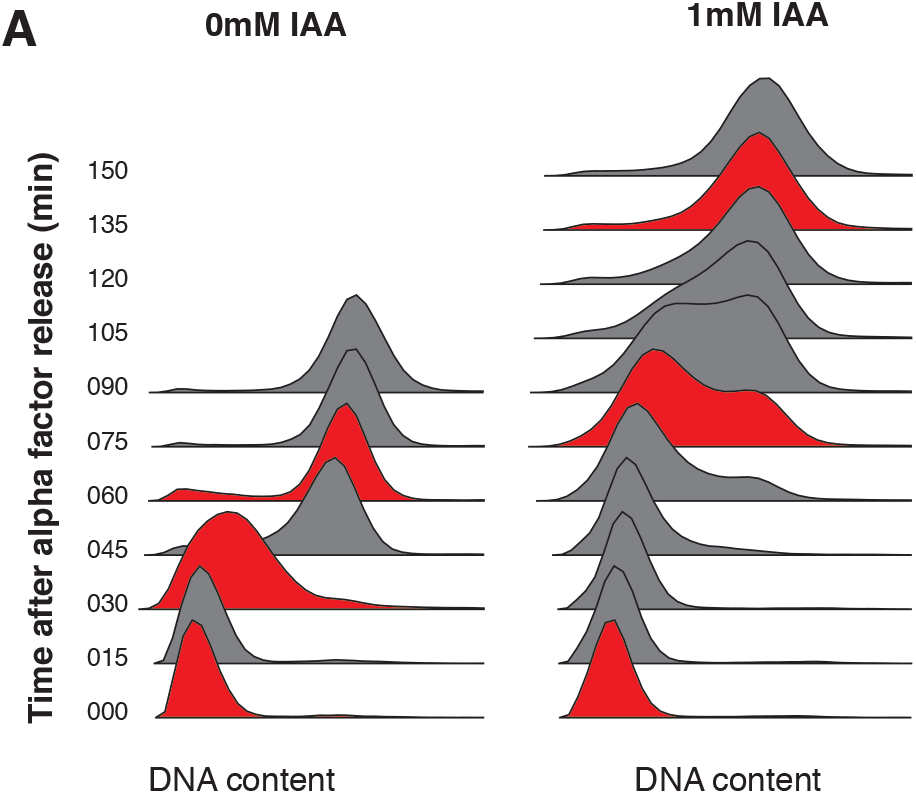
**A.** DNA content measured by flow cytometry for samples used for western blotting of Rfa1 in Fig. 5B. Red timepoints were chosen for G1, mid S, or late S samples.

